# Autoencoder Denoising for Network-Based Spatial Transcriptomics Data with Applications for Cell Signaling Estimation

**DOI:** 10.1101/2025.11.17.688924

**Authors:** Azka Javaid, H. Robert Frost

## Abstract

We propose an autoencoder-based framework for denoising networks estimated from Spatial Transcriptomics (ST) data for cell signaling analysis. Our method consists of an unsupervised encoder-decoder framework for denoising the network adjacency matrix and a supervised framework for cell signaling estimation. We validate our denoising component using the Frobenius norm metric for graphs simulated using the Barabasi–Albert (BA) and Erdős–Rényi (ER) models against reconstructions generated using the singular value decomposition (SVD). We then validate the cell signaling estimates generated using the supervised component on real ST data for the Wnt3-Fzd1 and Ephb1-Efnb3 interactions. We report that our framework achieves better adjacency matrix reconstructions for superlinear BA and dense ER graphs and generates cell signaling estimates that are both regionally specific and biologically plausible. An important contribution of this work is the application of neural networks for network-based cell signaling estimation using ST data and the benchmarking of autoencoder versus SVD denoising for different graph models.

## 1 Introduction

### 1.1 Overview of spatial transcriptomics technologies

Spatial transcriptomics (ST) technology enables estimation of mRNA abundance and quantification of spatial coordinates for thousands of genes and thousands to hundreds-of-thousands of locations [1]. ST technologies can broadly be categorized as imaging-based or sequencing-based. Imaging-based technologies function by imaging mRNAs in situ via microscopy and include in situ hybridization (ISH)-based methods such as MERFISH [2]. Sequencing-based technologies function by capturing tissue-specific mRNAs, synthesizing cDNAs and counting gene-specific sequences and include microdissection-based techniques such as Geo-seq [3] and positionally-barcoded, array-based methods such as Visium [1]. As compared to sequencing-based technologies, imaging-based techniques enable mRNA profiling at subcellular spatial resolution. In comparison, sequencing-based technologies can often profile larger tissue sections at a whole transcriptome-level [4]. While ST data can be represented using various mathematical and statistical structures, we particularly discuss network-based representations of ST data.

### 1.2 Network-based representation of spatial transcriptomics data

ST data is commonly mapped to a network model where nodes represent profiled spatial locations and edges capture transcriptomic and/or spatial distance. When both transcriptomic and spatial information is captured in the network, this enables estimation of gene expression heterogeneity and colocalization patterns that vary or decay over distance [5]. Estimation of distance-based colocalization patterns is especially appropriate for studying biological phenomena such as ligand-receptor-based cell signaling. Characterization of cell signaling is an important analysis objective for both scRNA-seq and ST data. Existing approaches for scRNA-seq data include CellPhoneDB [6] and CellChat [7], which perform estimation by leveraging expression information from intracellular agents down-stream or on the molecular pathway of a ligand-receptor interaction. Similarly, optimal transport-based methods like COMMOT [8] have been proposed to perform cell-cell communication for ST data by leveraging gene expression and location-specific information. Methods like Giotto [9] have leveraged network models of ST data represent spatial locations/cells using nodes with edges capturing spatial adjacency. While these methods characterize the signaling landscape at a granular, cell/spot-level, they are limited in that they do not directly incorporate spatial distance in the network construction. Another challenge with network-based representation of ST data is that it is limited by the noise and sparsity characteristic of ST assays. For ST data, the biological signal can be obscured by multiple factors including sparsity, technical artifacts and biological variability resulting from cell-type/phenotypic heterogeneity [10]. ST data quality is further impacted by coverage limitations in each sequencing unit and the maintenance of spatial positions during sequencing [11]. Given the noise inherent in the technologies used to generate ST data, there is a pressing need for effective denoising methods that can improve the accuracy of network-based ST models.

### 1.3 Singular value decomposition (SVD)-based reconstruction of adjacency matrices

We previously proposed our degree-centrality-based network analysis method for cell-cell communication analysis leveraging singular value decomposition (SVD)-based reconstruction of a ST-specific adjacency matrix [12]. In that method, we applied SVD on the normalized ST gene expression matrix. We then constructed a fully connected, directed and weighted adjacency matrix with edge weights set to the product of the reduced rank reconstructed expression for the ligand at the source location and receptor at the target location weighted by inverse squared distance between the locations. Following the adjacency matrix construction, we computed the weighted in-degree centrality on this adjacency matrix to estimate location-specific signaling activity of the target ligand-receptor interaction for the Wnt3-Fzd1, Ephb1-Efnb3 and Ptprc-Cd22 interactions. While the estimates generated by this approach are biologically plausible and regionally specific, a key limitation of our technique is that our SVD-based reconstruction is not capturing potential non-linearities in cell signaling profiles specific to a ligand-receptor interaction.

### 1.4 Autoencoder-based reconstruction of adjacency matrices

Autoencoder frameworks have been leveraged for spatial domain identification for ST data. STAGATE [13], for example, leverages graph attention-based autoencoder to integrate spatial information and gene expression profiles. Similarly, methods like SpaGCN [14] apply graph convolutional network to integrate gene expression and spatial data while techniques like SEDR [15] leverage variational graph autoencoder to embed spatial information with gene representations learned using deep autoencoder networks. A key limitation of these methods is that these techniques do not directly incorporate spatial distance within the network model to define gene expression-based edge weights that decay over distance.

In this manuscript, we describe a novel autoencoder-based framework for effective denoising of ST network models. For this task, we provide a benchmarking analysis that simulates graphs according to different structural and mechanistic properties to examine the accuracy of adjacency matrix reconstruction using either an autoencoder framework or SVD. This simulation study shows that our autoencoder framework achieves a more accurate network reconstruction as compared to an SVD-based approach when evaluated using the Frobenius norm. To demonstrate the practical utility of our autoencoder framework, we use it to denoise a network model of real ST data that is employed for network-based cell signaling estimation. Importantly, the autoencoder-based network denoising generates cell signaling estimates that are more regionally specific and biologically plausible.

## 2 Methods

### 2.1 Schematic

Our method proposes an autoencoder-based unsupervised component for denoising of the network adjacency matrix estimated from ST data and a supervised component for cell signaling analysis. This approach is visualized in Figure 1.

**Fig. 1.**
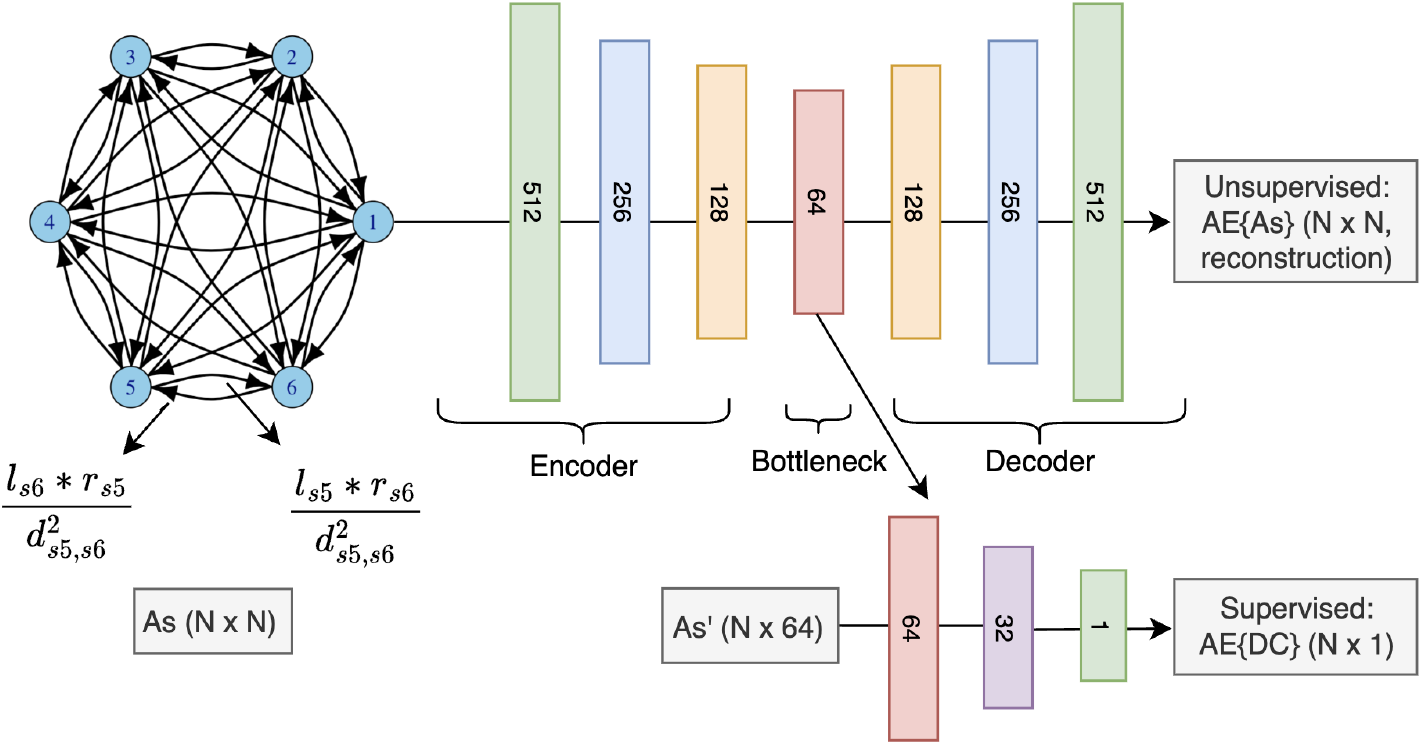
Unsupervised autoencoder(AE)-based denoising component for an input adjacency matrix *A*_*s*_ (*N × N* ) and a supervised component for cell signaling estimation using degree centrality (DC).

### 2.2 Data sources

#### Simulated data

To extensively validate the performance of the adjacency matrix reconstruction achieved using our autoencoder-based framework as compared to the reconstruction achieved using SVD, we simulated adjacency matrices according to different network models with random edge weights. Specifically, we simulated edge weights for a 3,000-by-3,000 adjacency matrix using the Barabási–Albert (BA) preferential attachment and Erdős–Rényi (ER) frameworks with the igraph package v2.1.4 in R. We simulated the BA-based graphs using the *sample pa* function with the n argument set to 3,000, the power argument (power of the preferential attachment with default of one, i.e., linear preferential attachment) varying from 0.1, 0.3, 1, 3 and 5 to explore a range of sublinear, linear and superlinear preferential attachment scenarios, and the m argument (number of edges to add in each time step) set to 10. We then simulated edge weights as standard uniform random variables to generate the ground-truth network. Lastly, we added Gaussian noise with mean 0 to our simulated ground-truth adjacency matrix. For evaluation purposes, we fixed the standard deviation of the Gaussian noise to 0.5. Similarly, we simulated the ER-based graph using the *erdos*.*renyi*.*game* function with the n argument (number of vertices) set to 3,000 and the p.or.m argument (probability for drawing an edge between two arbitrary vertices) varying between 0.3 and 0.9 to assess the influence of network density.

#### Real spatial transcriptomics data

To assess performance on real ST data, we applied our autoencoder framework to the posterior slice of a 10x Visium mouse brain ST dataset, consisting of 48,721 genes and 3,353 cells, which we accessed using the *LoadData* function from the SeuratData package v0.2.2.9001. We performed cell signaling estimation for the Wnt3-Fzd1 and Ephb1-Efnb3 interactions on this ST data since Wnt3 is implicated in processes including regulation of neurogenesis and repression of neural tumors [16, 17] and Ephb1 is essential for regulating accurate axon guidance as well as processes like cell migration and synapse formation [18].

### 2.3 Singular value decomposition-based denoising

We performed reduced rank reconstruction (RRR) on the *N* × *N* adjacency matrix using randomized SVD as implemented by the *rsvd* function from the rsvd package v1.0.5 [19]. Since we were interested in evaluating the SVD-based reconstruction against the autoencoder-based reconstruction of the input adjacency matrix under a range of aggressive and weak compression scenarios, we specified parameter *k* (the target rank for the low-rank decomposition) ranging from 5, 10 and 50.

### 2.4 Ligand-receptor network construction using real data

To perform cell signaling estimation for ST data, we constructed a fully connected, weighted and directed ligand-receptor activity network. We stored this network as an adjacency matrix A with nodes representing the individual tissue locations and edges representing weighted connections between two locations, weighted by the product of the normalized and reduced rank reconstructed lig- and and receptor gene expression scores. Lastly, we penalized this score by the inverse squared Euclidean distance between the two spots. We note that for the task of cell signaling estimation for real ST data, we computed reduced rank reconstruction using the *rsvd* function with the target rank set to 100. Following network construction, we computed the weighted in-degree centrality for each location by computing the sums of the weights over the columns. We report this degree centrality metric as the estimate of cell signaling activity for a given ligand-receptor pair at the target location. Further details about our ligand-receptor network construction and weighting mechanism can be found at [12].

### 2.5 Autoencoder architecture: unsupervised training component

Our autoencoder architecture (see Figure 1) consists of encoder and decoder components. While both the encoder and decoder frameworks are set as four-layer neural networks (512, 256, 128 and 64 units), the decoder reverses the latent embedding output from the encoder to reconstruct the original input adjacency matrix. For each layer, we set the activation function to the Rectified Linear Unit (ReLU), with the exception of the final layer where the activation function is set to sigmoid. We set the activation function to ReLU for most layers since it is computationally efficient and prevents the problem of vanishing gradients. In comparison, we specified sigmoid as the activation function in the final layer since it ensures that reconstructed values are on a similar scale as the input data. For each layer implementing the ReLU activation function, we specified the kernel initializer to the He uniform [20]. In comparison, for the final layer implementing the sigmoid activation function, we specified the kernel initializer to the Glorot uniform [21]. In addition to the activation function specification, we specified the regularizer to L2 with regularization factor of 0.001 in every layer except the output layer. To further ensure that our reconstruction was generalizable and not overfitting, we specified layer-specific dropout rates successively ranging from 0.3, 0.4, 0.5 and 0.6 in the layers with 512, 256, 128 and 64 units, respectively. Lastly, we specified batch normalization between all intermediate layers to ensure adequate training stabilization.

### 2.6 Autoencoder architecture: supervised training component

To validate that our autoencoder learns meaningful structural representations, we implemented a supervised component that predicts degree centrality using the 64-dimensional latent embeddings from the autoencoder’s bottleneck layer. This three-layer neural network (64, 32 and 1 units) implements ReLU activation for hidden layers and linear activation for the output layer, with He uniform kernel initialization [20]. We further applied batch normalization after the first layer and dropout regularization (0.3 and 0.2 rates) to prevent overfitting. This model uses Adam optimization with gradient clipping and early stopping and trains on the z-score normalized degree centrality target scores computed on the fully connected, weighted and directed ligand-receptor activity networks for real ST data. The supervised component of our autoencoder-based framework ensures that the unsupervised autoencoder representations capture essential network topological information and cell signaling distributions beyond those captured by the weighted in-degree centrality scores.

### 2.7 Benchmarking

We assessed the accuracy of the reconstructions achieved using our autoencoder-based framework and SVD using the Frobenius norm. For our task, we computed the Frobenius norm between our simulated ground-truth adjacency matrix (*A*_true_) (i.e., before the noise addition) and each of the reconstructions achieved using the autoencoder-based framework (*A*_*AE*_) and SVD (*A*_*SV D*_) on the input matrix (i.e., the simulated ground-truth adjacency matrix and noise: *A*_*s*_) and between *A*_true_ and *A*_*s*_. We evaluated the reconstruction error using the Frobenius norm metric as defined.

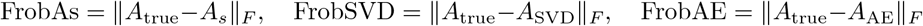

## 3 Results

### 3.1 Autoencoder-based framework achieves better reconstruction loss for superlinear Barabási–Albert graphs

We first examine variations in the Frobenius norm metric for matrix A generated using the BA model. For this task, we explore the influence of the power attribute on the computed metrics by holding the number of edges to be added at each time step constant at 10 and noise simulated using a fixed standard deviation estimate of 0.5. As shown by Table 1, as power increases, performance of our autoencoder-based reconstruction improves as indicated by lower Frobenius norm as compared to the SVD-based reconstruction. For example, when reconstructed using our autoencoder-based framework, the Frobenius norm decreases from 99.69 to 63.93 (*Δ* = 35.76) for graphs constructed with power = 1 versus power = 3. This reduction is larger than that observed with the SVD-based reconstruction, where the Frobenius norm decreases from 93.52 to 71.26 (*Δ* = 22.26). Additionally, the difference in the Frobenius norm computed for the autoencoder-based reconstruction relative to the SVD-based reconstruction decreases as power increases from 1 (*Δ* = −7.33 for power = 3, *Δ* = −9.78 for power = 5). Since power *>*1 corresponds to a superlinear attachment model, where a small number of vertices become disproportionately highly connected, a phenomenon sometimes described as an amplified form of the “rich-get-richer” mechanism [22], our results indicate that the autoencoder-based framework is better able to capture the nonlinear growth patterns implied by this graph construction.

**Table 1.** Frobenius norm (Frob) computation for matrix *A* generated using the Barabási–Albert (BA) model with the power attribute varying from 0.1, 0.3, 1, 3 and 5. The number of edges added at each time step is fixed at 10, with edge weights drawn from a random uniform distribution and noise generated from a normal distribution (mean = 0, standard deviation = 0.5). RRR is performed using SVD with target rank set to 5. Bold text format is used to indicate method with lowest Frobenius norm for each experiment.

| Method | PA(power=0.1) | PA(power=0.3) | PA(power=1) | PA(power=3) | PA(power=5) |
| --- | --- | --- | --- | --- | --- |
| FrobAs | **86.66** | **86.73** | **86.47** | 86.07 | 86.07 |
| FrobSVD | 99.22 | 99.34 | 93.52 | 71.26 | 71.25 |
| FrobAE | 104.72 | 106.09 | 99.69 | **63.93** | **61.47** |

We next examined variations in the Frobenius norm metric for matrix A generated using the BA model with power fixed to be 3, the number of edges to be added at each time step held constant at 10 and noise simulated using a fixed standard deviation estimate of 0.5. We then varied the target rank k to better explore the influence of an aggressive (i.e., k = 5) versus weak (i.e., k = 50) compression on the loss computed using the reconstruction generated with SVD. As Table 2 shows, the Frobenius norm metric increases as rank is changed from 5 (Frobenius norm = 71.26) to 10 (Frobenius norm = 86.08) and 50 (Frobenius norm = 86.07) for reconstructions generated using the SVD. Our findings indicate that higher-rank SVD reconstructions may suffer from over-fitting to noise in the adjacency matrix representation. We further hypothesize that BA networks, especially when generated using power = 3, likely have a low effective rank structure, where most topological information is concentrated in a small number of dominant singular vectors. Therefore SVD-based performance naturally degrades as rank exceeds a threshold due to potential amplified noise.

**Table 2.** Frobenius norm (Frob) computation for matrix *A* generated using the Barabási–Albert (BA) model with the power attribute fixed at 3. The number of edges added at each time step is fixed at 10, with edge weights drawn from a random uniform distribution and noise generated from a normal distribution (mean = 0, standard deviation = 0.5). RRR is performed using SVD with target rank (*k*) varying from 5, 10, and 50. Bold text format is used to indicate method with lowest Frobenius norm for each experiment.

| Method | PA($k = 5$) | PA($k = 10$) | PA($k = 50$) |
| --- | --- | --- | --- |
| FrobAs | 86.07 | 86.07 | 86.07 |
| FrobSVD | 71.26 | 86.08 | 86.07 |
| FrobAE | **63.93** | **63.55** | **63.55** |

### 3.2 Autoencoder-based framework achieves better reconstruction loss for dense Erdős–Rényi graphs

Following the assessment of the Frobenius norm metric for matrix A generated using the BA model, we next assessed these metrics for matrix A generated using the ER model. For this task, we varied the edge probability from 0.3 and 0.9, fixed the standard deviation used to simulate noise to 0.5 and varied the rank k for SVD from 5, 10 and 50. We first observe that the Frobenius norm for the reconstruction generated using our autoencoder-based framework decreases relative to the SVD-based reconstruction as the edge probability increases from 0.3 to 0.9 (see Table 3). For example, when reconstructed using our autoencoder-based framework, the Frobenius norm for edge probability of 0.3 is 835.7 as compared to 834.9 for the reconstruction constructed using a rank of 5 with SVD. While the Frobenius norm for edge probability of 0.3 for the reconstruction built using our autoencoder-based framework increases to 937.2 for network constructed using edge probability of 0.9, the difference in the Frobenius norm relative to the SVD-based reconstruction (Frobenius norm = 939.8) decreases, which is especially noticeable as the rank increases from 5 to 10 and 50 (*Δ* = −2.6 for rank 5, *Δ* = −5.9 for rank 10, *Δ* = −31 for rank 50). Our findings indicate that as the edge probability increases from 0.3 to 0.9 in the ER model, the autoencoder-based framework is able to better capture the dense network structure as compared to the SVD-based reconstruction.

**Table 3.**
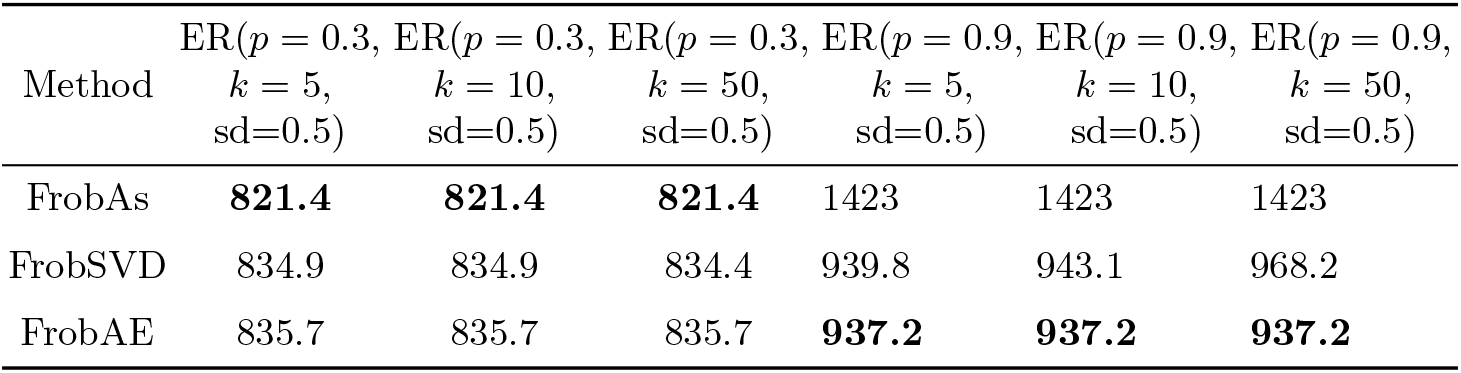
Frobenius norm (Frob) computation for matrix *A* generated using the Erdős–Rényi (ER) model with edge probability *p* set to 0.3 or 0.9. Edge weights are drawn from a random uniform distribution and noise is generated from a normal distribution with mean 0 and standard deviation (sd) fixed at 0.5. RRR is performed using SVD with target rank (*k*) varying from 5, 10, and 50. Bold text format is used to indicate method with lowest Frobenius norm for each experiment.

| Method | ER($p = 0.3$,<br>$k = 5$,<br>sd=0.5) | ER($p = 0.3$,<br>$k = 10$,<br>sd=0.5) | ER($p = 0.3$,<br>$k = 50$,<br>sd=0.5) | ER($p = 0.9$,<br>$k = 5$,<br>sd=0.5) | ER($p = 0.9$,<br>$k = 10$,<br>sd=0.5) | ER($p = 0.9$,<br>$k = 50$,<br>sd=0.5) |
| --- | --- | --- | --- | --- | --- | --- |
| FrobAs | **821.4** | **821.4** | **821.4** | 1423 | 1423 | 1423 |
| FrobSVD | 834.9 | 834.9 | 834.4 | 939.8 | 943.1 | 968.2 |
| FrobAE | 835.7 | 835.7 | 835.7 | **937.2** | **937.2** | **937.2** |

### 3.3 Autoencoder-based framework generates regionally specific cell signaling estimates

We next applied our autoencoder-based framework to generate cell signaling estimates for the Wnt3-Fzd1 and the Ephb1-Efnb3 interactions in mouse brain ST data. We visualize the cell signaling estimates generated using our autoencoder-based framework against the cell signaling estimates generated using our weighted in-degree centrality method [12].

As Figure 2 shows, the cell signaling profiles generated using our autoencoder-based framework (AE_trainedDC) for the Wnt3-Fzd1 interaction are more regionally specific to the mouse cerebellum region as compared to the cell signaling profiles generated using the in-degree centrality method (targetDC), which show more expression heterogeneity. Specifically, the signaling profiles generated using the in-degree centrality method are mostly specific to both the cerebellum and the thamalus regions while the profiles generated by our autoencoder-based framework are highly specific to the cerebellum region. Similarly, the cell signaling profiles generated by our autoencoder-based framework for the Ephb1-Efnb3 interaction show clear specificity to the hindbrain region (i.e., regions of the cerebellum, the medulla oblongata and the pons). In comparison, the profiles generated by the in-degree centrality method are noisier and lack clear specificity to the hindbrain region. Our signaling profiles generated for the Wnt3-Fzd1 and Ephb1-Efnb3 interactions have biological plausibility as Wnt3 is typically expressed in the mouse cerebellum region during postnatal development [17] while Ephb1 is known to be involved in hindbrain tangential cell migration [23].

**Fig. 2.**
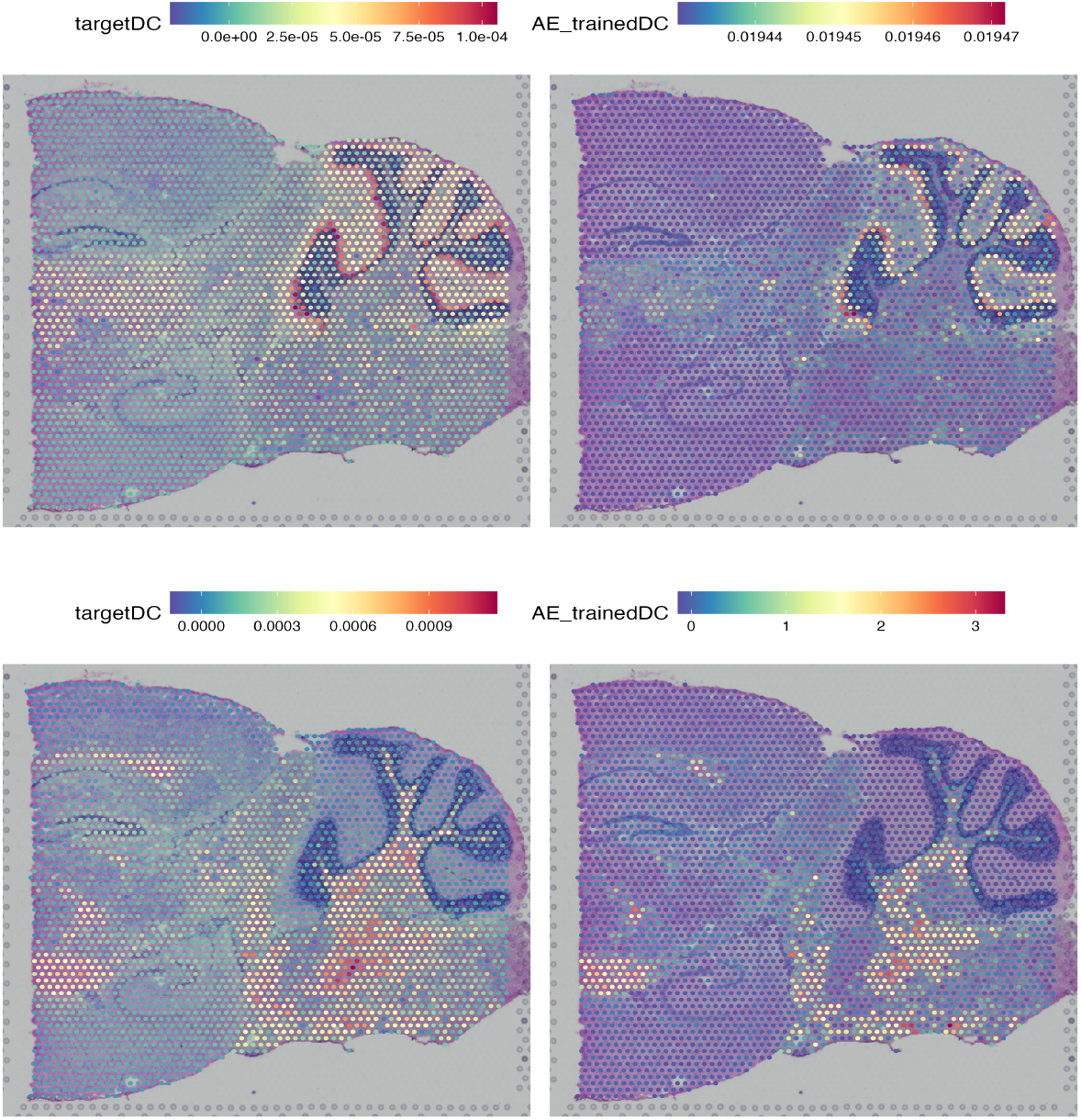
Cell signaling estimates generated using the weighted in-degree centrality method (trainedDC) and the autoencoder-based framework (AE_trainedDC) for the Wnt3-Fzd1 interaction (top) and for the Ephb1-Efnb3 interaction (bottom).

## 4 Conclusions and future directions

In this paper, we propose a novel autoencoder-based framework for denoising of network-based adjacency matrix data and cell signaling estimation of ST data. We validated our method by computing the reconstruction loss using the Frobenius norm for graphs simulated using the Barabási–Albert (BA) and Erdős–Rényi (ER) models. We also validated the cell signaling estimates generated by our method on real ST data for the Wnt3-Fzd1 and Ephb1-Efnb3 interactions. We report that our autoencoder-based framework achieves better reconstruction loss for superlinear BA and dense ER graphs and generates cell signaling estimates that are more regionally specific and biologically plausible than comparative techniques. For future extensions of this paper, we plan to expand our framework to multiple interactions and graph models with different parameter settings.

